# Learning vs. minding: How subjective costs can mask motor learning

**DOI:** 10.1101/2022.02.11.479978

**Authors:** Chadwick M. Healy, Max Berniker, Alaa A. Ahmed

## Abstract

When learning new movements some people make larger kinematic errors than others, interpreted as a reduction in motor-learning ability. Consider a learning task where error-cancelling strategies incur higher effort costs, specifically where subjects reach to targets in a force field. Concluding that those with greater error have learned less has a critical assumption: everyone uses the same error-canceling strategy. Alternatively, it could be that those with greater error may be choosing to sacrifice error reduction in favor of a lower effort movement. Here, we test this hypothesis in a dataset that includes both younger and older adults, where older adults exhibited greater kinematic errors. Utilizing the framework of optimal control theory, we infer subjective costs (i.e., strategies) and internal model accuracy (i.e., proportion of the novel dynamics learned) by fitting a model to each population’s trajectory data. Our results demonstrate trajectories are defined by a combination of the amount learned and strategic differences represented by relative cost weights. Based on the model fits, younger adults could have learned between 65-90% of the novel dynamics. Critically, older adults could have learned between 60-85%. Each model fit produces trajectories that match the experimentally observed data, where a lower proportion learned in the model is compensated for by increasing costs on kinematic errors relative to effort. This suggests older and younger adults could be learning to the same extent, but older adults have a higher relative cost on effort compared to younger adults. These results call into question the proposition that older adults learn less than younger adults and provide a potential explanation for the equivocal findings in the literature. Importantly, our findings suggest that the metrics commonly used to probe motor learning paint an incomplete picture, and that to accurately quantify the learning process the subjective costs of movements should be considered.

**Author Summary:** Here we show that how a person values effort versus error in their movements has an impact on their overall strategy for performing those movements and adapting to a novel environment. When error alone is considered as a measure of learning, it appears that certain populations such as older adults are significantly worse at learning new motor tasks. However, using an optimal control framework, we are able to parse out differences in how much a population or person has learned, as well as how they subjectively value factors such as effort and error. In the case of older adults, we show that they could be learning as much as younger adults but exhibit larger errors because they care more about expending extra effort to reduce them.

## Introduction

When people are introduced to a novel environment, they initially experience large errors relative to their intended performance. Through repeated practice, they systematically reduce these errors, learning how best to perform the task [1–10]. Because the amount a person has learned is not directly assessable, we are reliant on indirect measures such as error that reflect the latent state of how much a person has learned. However, error-reduction in and of itself does not necessarily equate to learning a model of the task. First, numerous studies have shown that a model-free process, such as impedance control, can cancel error without the necessity of learning an accurate model of the task [11,12]. Second, even when assuming similar model-free approaches, when comparing across subjects or populations, variation in strategies must be considered. In this study, we focus on the latter point. We ask whether greater error may not be a result of learning less, but rather, a strategic trade-off between movement errors and other similarly important factors.

Movement decisions and learning are not solely driven by error reduction. People optimize for other factors such as reward, effort, time, and risk. When people move faster, they move with greater error [13] and tend to have trajectories which minimize the variance of the end or target position [14]. Yet, people still choose to move faster in the presence of greater reward in both arm reaches [15] and saccades [16]. Reward is discounted by time, thus people and animals are willing to expend more effort to achieve an earlier reward [17–19]. In addition to error, studies have shown that effort is reduced through the learning process [4,8], suggesting a simultaneous optimization of both these factors [20]. Studies have also shown learning rate is increased by increasing rewards [5], increasing consequence of an error [21], or by manipulating the level of uncertainty of state or the environment [22,23]. Traditionally interpreted as learning to a lower extent, increased residual error in an adaptation task is influenced by the consequences of an error [24] or the level of effort [20]. These studies, in conglomeration, show that there is more at play in motor learning than just error reduction. When evaluating the extent of learning in motor learning studies, residual errors do not necessarily mean a reduced amount of learning has occurred, but rather, subjects could be optimizing for factors in addition to error.

Optimal feedback control theory offers a formalization of these trade-offs and has been used to describe human movements [25–28]. Using this framework, subjective costs are quantified and used to develop a control policy, offering insight beyond kinematics into the subjective strategies and decisions of each individual. Izawa and colleagues showed that in an adaptation task, higher error is acceptable, and indeed optimal, if the total movement cost includes not only error but effort as well [28]. Others have used an optimal control model to formalize trade-offs between reward and effort that could predict not only which movement would be made, but also how that movement would be performed [19]. Optimal control models offer insight that is deeper than a comparison of *how* the kinematics of two movements differ; by investigating the subjective costs used, these models offer an explanation for *why* the movements differ.

Many motor learning studies implicitly assume populations use identical strategies when learning new motor tasks, thus kinematic differences between populations are evidence for a difference in the ability to learn. Using error as a correlate for the amount learned, prior literature shows mixed results as to whether motor learning capability declines with age. A number of studies have concluded that our ability to learn new motor tasks declines with age [29–34], attributing this observation to underlying causes such as motor variability [35,36], sensory deficits [37,38] and attention [39]. However, the causes are inconsistent and are highly dependent on the task structure, complexity, and familiarity [40]. In some cases, older adults learn to the same extent as younger adults [41–45]. In other studies, we see older adults exhibit different movement strategies than younger adults [45,46]. One study has shown that older adults put higher subjective value on effort than their younger counterparts [47]. The observed kinematic differences between older and younger adults [45], combined with their difference in strategies and subjective cost values, suggest that it is important to isolate and investigate these variables separately.

Specific to reach adaptations to a curl field, a few recent studies have found no differences in learning between older and younger adults [41,43,44], however each use slightly different protocols. Comparing protocols to the study conducted by Huang and Ahmed [29], which shows significant differences, protocols differed in curl field gains, reach distances, reach directions, and/or inter-trial intervals. These differences in protocol may be the reason for why differences are observed in Huang and Ahmed where others have not: the task demands more effort which may have accentuated the effect of age-related changes in sensitivity to effort. Huang and Ahmed’s protocol uses a higher value for curl gain (−20 N-s/m), longer reaches (20 cm), and only brief pauses (no robot-guided reset to the starting point, reaches starting every two seconds).

Here we ask whether observed age-related errors during motor learning in a velocity-dependent force field can be explained by a difference in subjective costs rather than a difference in learning. We focus specifically on a dynamics learning task, wherein subjects must reach in a velocity-dependent force field. In this task, learning involves a trade-off between effort and error reduction. Some recent studies have shown that older and younger adults perform similarly in adapting to a velocity dependent force field. However, Huang and Ahmed observe a difference between age groups in their 2014 study [29]. We use data from this previously published study and fit an optimal feedback control model to the younger and older adult reach trajectories. Using a simple model, we first demonstrate how differences in strategies, which we quantify through subjective costs, can give rise to changes in learned behavior usually interpreted as reduced learning. Next, we determine the range of strategies that can explain the observed behavior. Together, the results demonstrate a large overlap across the younger and older adults in the proportion learned if their subjective costs are different. In particular, the two groups appear to learn similar amounts, but older adults place a higher subjective cost on effort required to reduce kinematic errors.

## Results

### Modeling subjective cost trade-offs

Trade-offs between subjective costs and the amount of learning can produce equivalent control laws, and thus trajectories. This concept is best illustrated with a simple model. Let us consider a one-dimensional linear dynamical system:

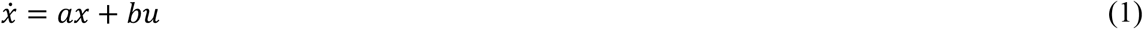

where *x* represents the state, *u,* the control, and *a* and *b* parameterize the dynamics. We can model learning as the process of estimating an internal model of the state dynamics where *â* is the estimate of the true value *a*, scaled by *ε,* the “proportion learned”.

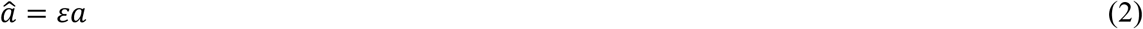

Given this internal model, we define the conventional linear quadratic cost, *J,* which penalizes state deviations from zero (kinematic error) and control (effort) with weights *q* and *r,* respectively.

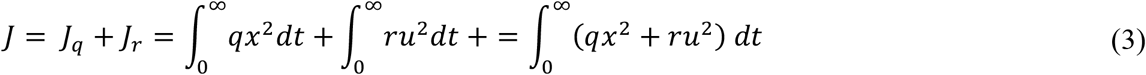

The solution to this cost is the well-known LQR solution, and the control, *u*, is expressed below:

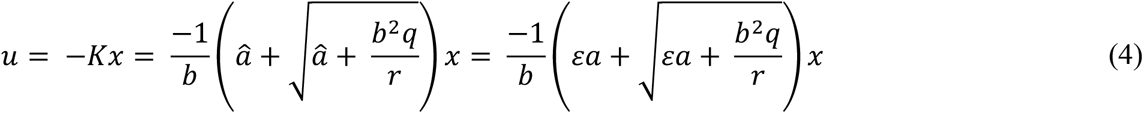

From this equation we see that similar control laws and resulting trajectories can be created through different combinations of the parameters, *q, r,* and *ε.* To illustrate, the controller obtained for one particular value of *q*, *r*, and *ε* can be identical to another controller with a reduced *ε* by increasing the penalty on control input, *r*, or decreasing the penalty on state, *q*, or a combination of the two. Accordingly, two different trajectories do not necessarily indicate a different internal model of the dynamics; this could be a result of having differing costs. In other words, there is no unique mapping from the control law and trajectory to the proportion of the dynamics learned. These trade-offs are visualized in Fig 1. In the case of this simple model, *q* and *r* represent the subjective costs of a person or population and *ε* represents the proportion of the dynamics that person or population has learned. This relationship clearly demonstrates how changes in subjective costs across individuals can mask differences in how much they have learned.

**Fig 1.**
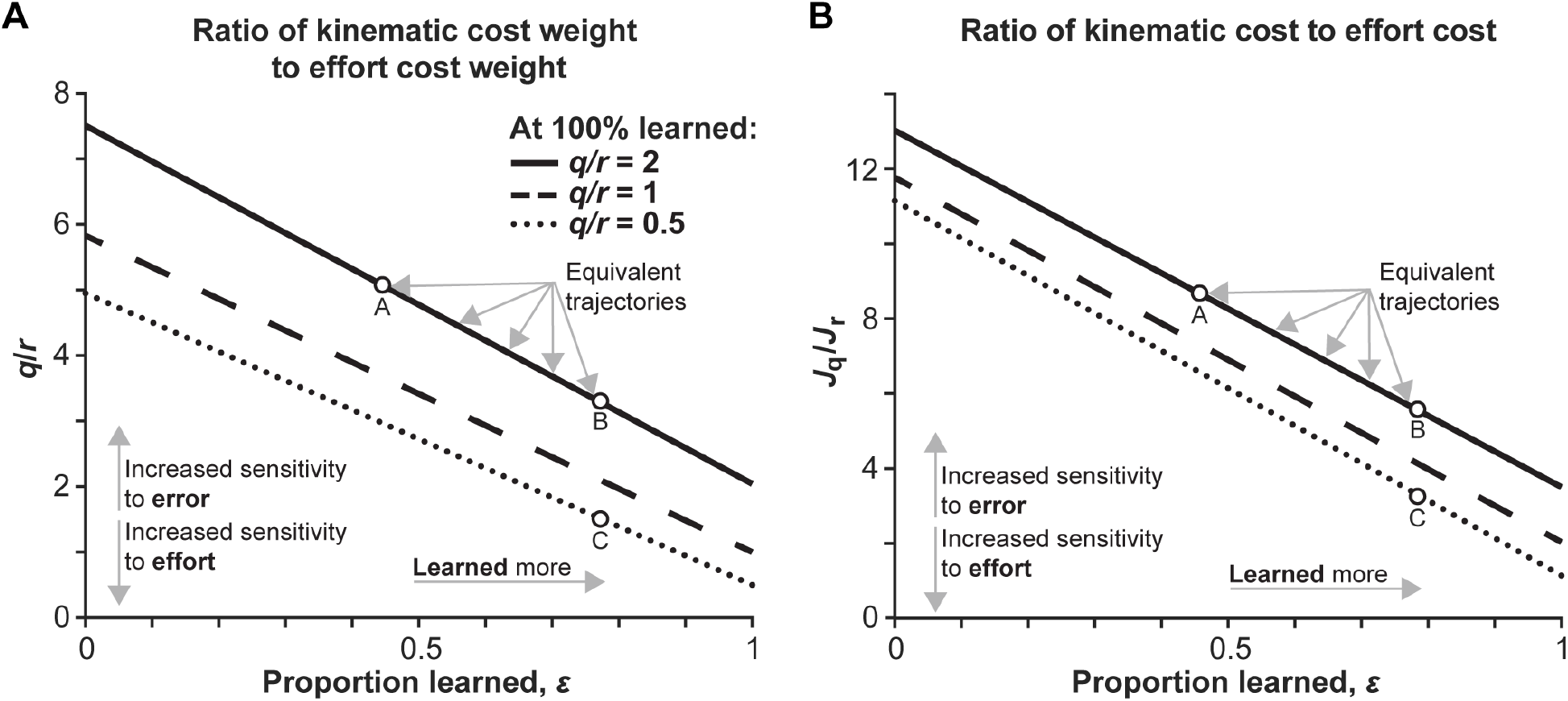
Trade-offs between subjective costs and proportion learned can produce equivalent trajectories. Each line represents a unique solution produced by ratios of kinematic cost weight to effort cost weight of 2, 1 and 0.5 with perfect knowledge of the dynamics. As the extent of learning is reduced, moving left along the x-axis, we see that in order to produce an equivalent trajectory, represented by each line, the ratio of kinematic error costs to effort costs would need to increase. (A) shows the ratio of specific cost weights, *q* and *r;* (B) shows the ratio of the total kinematic cost (*J_q_*) combining *q* and *x* to the total effort cost combining *r* and *u* (*J_r_*). Both (A) and (B) show that a trajectory with a lower proportion learned, but with a ratio of kinematic costs to effort costs is greater could be identical to a trajectory with a higher proportion learned but with a lower ratio of kinematics costs to effort costs. Conversely, if two groups used the same strategy (ratio of cost weights) and produced different trajectories, then they must have learned a different amount. For example, Point A and B are both equivalent trajectories because they fall along the same line, however, Point A has learned less than Point B. Thus, to produce the same trajectory as Point B, the ratio of kinematic costs to effort costs is higher for Point A. Next, Point B and Point C have learned to the same extent, but because Point B has a higher relative kinematic cost to effort cost than Point C, they produce different trajectories (i.e., they are not on the same line). These two comparisons show that kinematic observations alone do not correlate to a specific proportion learned.

### Differences in reaching behavior between older and younger adults

Using the example illustrated above, we extend the model to apply it to an experimental dataset of subjects performing arm reaches in a motor learning task [29]. In that study, younger and older adults made planar arm reaches while holding onto the handle of a robotic manipulandum (shown in Fig 2). After a baseline period of 200 reaching movements between a start position and target circle, the robot applied a velocity-dependent force to the hand, acting perpendicular to its velocity by the following equation:

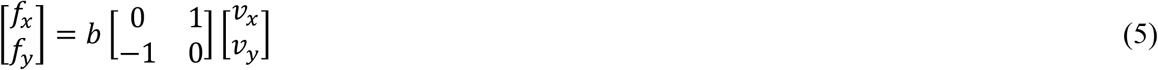

**Fig 2.**
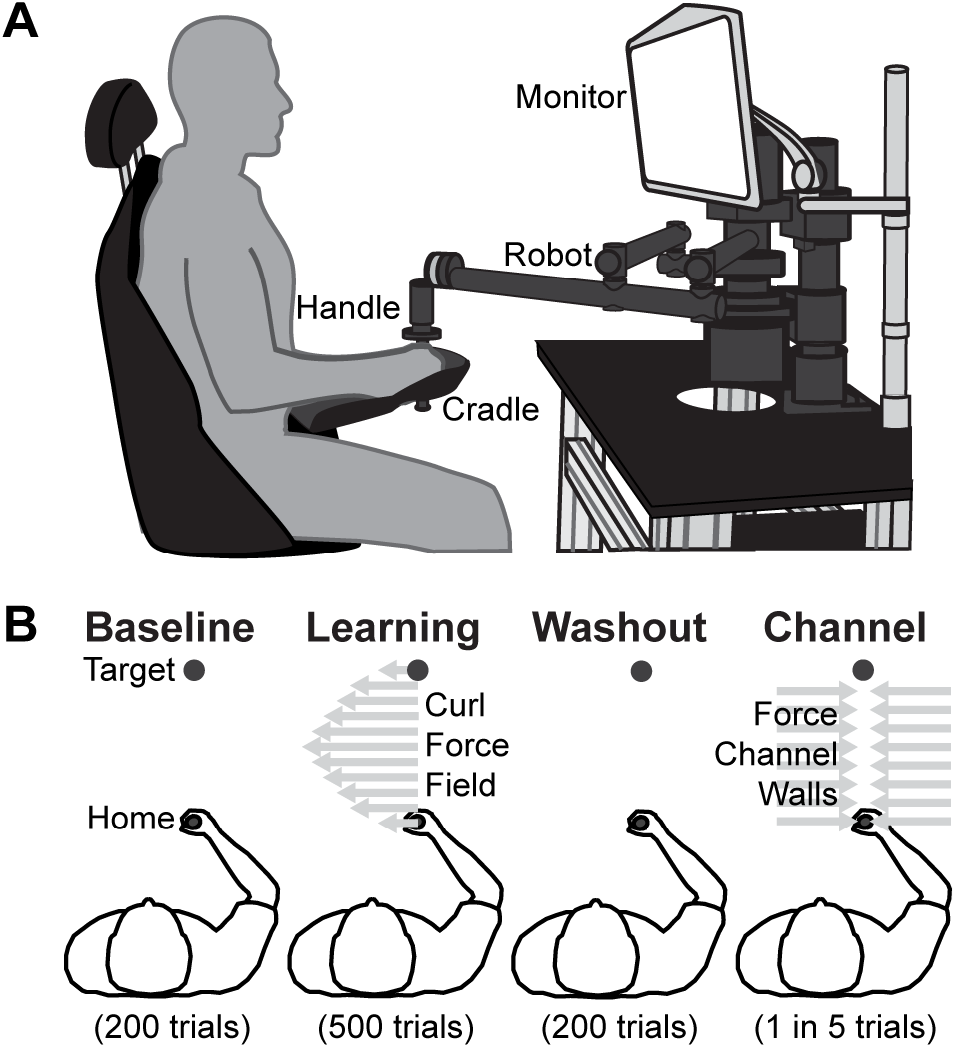
Experimental setup and conditions. (A) Subjects controlled a cursor on a computer screen by making reaches while grasping a robotic manipulandum, reaching anteriorly from one circle to a single target circle. (B) Subjects experienced 900 trials of varying dynamics. *Baseline* and *washout* phases have no perturbations, while *learning* imparted a curl field force specified in Equation 5 and represented by the gray arrows pushing them perpendicular to the direction of their reach. *Channel trials* were interspersed throughout the experiment, where subjects moved along a straight-line path to the target within a force channel. These trials are used to estimate the extent to which subjects were compensating for the dynamics by measuring the force that subjects pushed into the force channel wall, represented in gray. Details of how the channel trial is implemented are summarized in the Methods.

In Equation 5, *b* is the curl field gain, which was held constant at −20 N-s/m during the learning phase. As is typical for this paradigm, the force perturbation led to large deviations (i.e., errors) from the baseline movements that were subsequently reduced over many reaches. Abrupt removal of the force field led to large deviations in the opposite direction. To measure the horizontal force exerted, subjects were exposed to a force channel trial every five trials throughout the experiment, where reaches appeared to travel in a straight-line path towards the target. While the trends in error onset and reduction were similar in both younger and older adults, there were distinct differences in how they chose to reach the target.

Throughout our analyses, we quantify performance using three common metrics of learning: maximum perpendicular error, maximum perpendicular force, and a trajectory-derived adaptation index. Maximum perpendicular error is measured in non-channel trials and is calculated as horizontal deviation from a straight-line trajectory from the start position to the target. Maximum perpendicular force is measured in channel trials and is a coarse reflection of learning as it is a measure of the anticipatory force the subject is generating to counter the force field. The adaptation index normalizes the time profiles of anticipatory force by the velocity of the movement, purportedly correlating to a subject’s estimate of the curl field gain. However, this metric is a reflection of desired error cancellation: if force is perfectly correlated to velocity in a curl field, then reaches will have no horizontal deviation. As shown by Izawa and colleagues [28], some horizontal deviation is optimal even with perfect adaptation, thus adaptation index is influenced by subjective strategies. We focus on performance at four phases of the experiment: the last ten trials of the baseline period (late baseline), the first five and last ten trials in the learning period (early learning and late learning), and the first five trials after the perturbation was removed (early washout). To increase confidence by including more data, ten trials were included during the late phases, as these were less variable trial to trial. Because adaptation and de-adaptation occur so quickly, only the first five trials were considered during early learning and early washout.

Younger adults exhibited smaller perpendicular position errors than older adults in late learning (−0.553±0.169 cm versus −1.20±0.185 cm, *t*(26) = 2.53, *p*= 0.0176). Younger adults also exhibited greater maximum perpendicular force at late learning (11.4±1.45 N versus 8.06.±1.15 N, *t*(26) = 1.82, *p* = 0.0797), and a higher adaptation index at late learning (0.851±0.0398 versus 0.693±0.0510, *t*(26) = 2.38, *p* = 0.0248). Collectively, these metrics have been interpreted as older adults learning less than younger adults [29]. However, this conclusion does not consider the potential strategic differences between older and younger adults that may also cause these observed differences.

### Validating the arm reach model

To describe these trajectories, we model the limb as a point mass that moves in a two-dimensional plane. Similar to the simple model described above, we assume an internal model of the curl force field is parameterized by the gain. The model’s foundation is adapted from Izawa et al. 2008, where this internal model of the state dynamics (what we call “proportion learned”) was used to calculate the control law [28], however, our model assumes that the state is deterministic and perfectly observed. The model also includes higher derivative terms analogous to muscle activation filters [27]. The model uses a linear dynamical system, which includes hand position, velocity, force, rate of force, and target position as state variables. The model uses a cost function to determine its control law, where the cost function includes kinematic costs that penalize hand position error from target and horizontal velocity, effort-based cost that penalize hand force, rate of change of hand force, second derivative of hand force (control input), and target accuracy costs that penalize terminal position error, terminal velocity, terminal force, and terminal rate of change of force. Throughout this study, we represent subjective strategy as the log transform of the ratio of the sum of kinematic costs to effort costs, frequently shortened to “cost ratio”.

To investigate model behavior and predict possible findings between older and younger adults, we manually manipulated costs and proportions of learning and inspected the resulting trajectories run through the forward model (Fig 3A). As the cost ratio of error to effort increases, meaning the subject would care more about reducing error than expending effort, we see reduced horizontal deviations at low levels of learning in the curl field. At high amounts of learning, with low error-to-effort costs, we see trajectories with horizontal deviations into the curl field as predicted by Izawa and colleagues [28]. With a high cost ratio and high amounts of learning, we see straight trajectories that do not deviate horizontally. During washout, we see the same trend: at high amounts of learning, lower cost ratios deviate more than higher cost ratios. These trends present in the trajectory matches expectations, and shows the model is producing the expected results from our manual manipulations.

**Fig 3.**
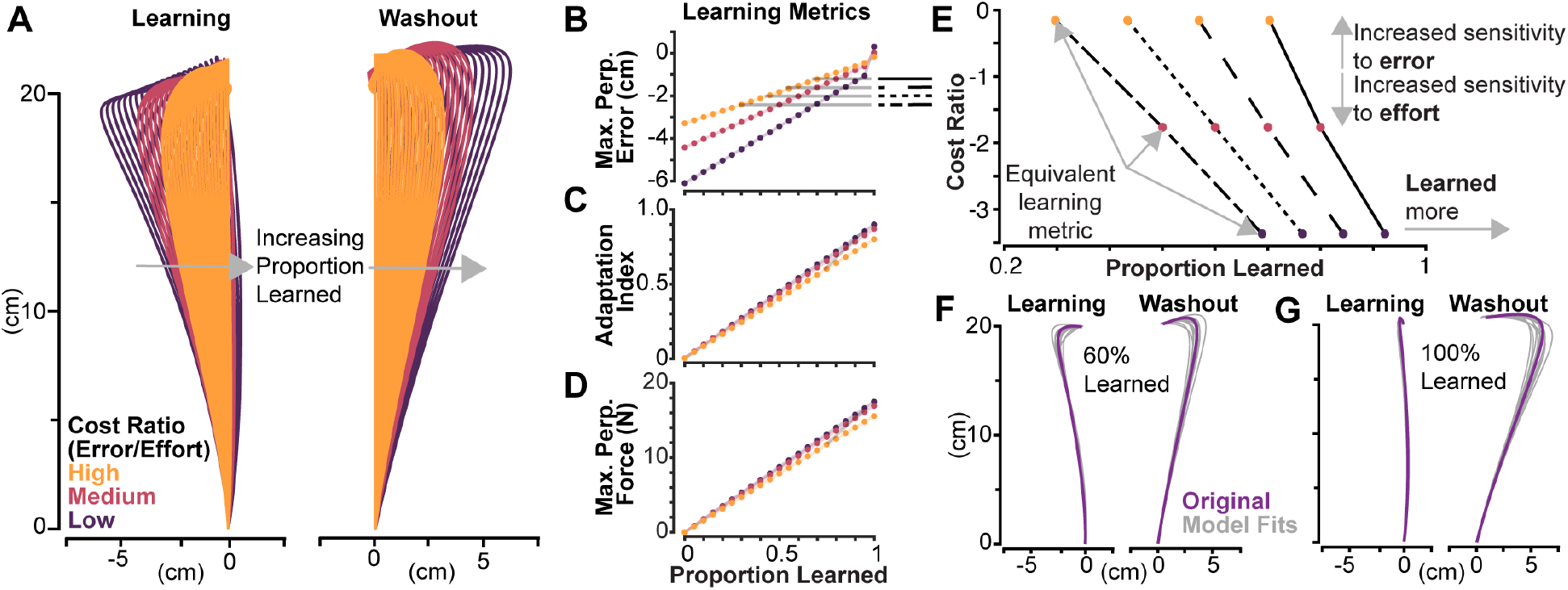
Model based predictions and validation. (A) Trajectories in the curl field, i.e., “Learning” and null field “Washout” with various proportion learned are shown for three types of cost ratios: “high”, “medium” and “low” in yellow, red, and purple, respectively. The corresponding learning metrics maximum perpendicular error, adaptation index, and maximum perpendicular for combinations of cost ratios and proportion learned are shown in (B)-(D). Lines connecting trajectories with equivalent maximum perpendicular are shown in (E). These lines correspond to the gray horizontal lines in (B), where the solid black line corresponds to a proportion learned of 80% for the medium cost ratio, and the dashed, dotted, and dash-dot to 70%, 60%, and 50%, respectively. The same learning metric can be produced by increasing sensitivity to error and a lower proportion learned, or vice versa, showing the same trend predicted by the simple model. Trajectories in (F) and (G) show model fits in gray relative to the original trajectory in purple for 60% and 100% proportion learned in the low-cost ratio condition. These results were used to validate the search function’s ability to back out cost ratios and proportion learned.

Fig 3B-D visualizes the trends in learning metrics for trajectories in the learning phase across cost ratios. As the proportion learned increases, we see maximum perpendicular error steadily decrease but go beyond zero in lower cost ratios. Additionally, we see adaptation index and maximum perpendicular force increase as learning increases. For this particular set of cost ratios, lower ratios have a slightly greater slope than higher cost ratios in Fig 3C and 3D, though this trend is not consistent across all combinations of costs. In some cases, the opposite can be true, where higher cost ratios have a greater slope. Clearly shown in Fig 3B-D, evidenced by their differing slopes, a given learning metric value is not uniquely specified by a proportion learned. Instead, this value is a function of both cost ratio and proportion learned.

Using the generated sets of trajectories shown in Fig 3A, we observe the same predictions made in the simple model and Fig 1: equivalent trajectories can be produced by decreasing cost ratio and increasing learning, or vice versa. In Fig 3E, values of 50%, 60%, 70%, and 80% proportion learned for the medium cost ratio were taken as baselines, then we find proportion learned values that produced equivalent maximum perpendicular error values in the high and low cost ratios. Values for the cost ratio were calculated by taking the log transform of the sum of the kinematic or “error” costs divided by the effort costs. The less negative the value (e.g., a “high” cost ratio), the more sensitive to error. The solutions show consistent negative slopes, where the high cost ratio solution produces an equivalent maximum perpendicular error with a lower proportion learned, and the lower cost ratio could produce the same maximum perpendicular error with a higher proportion learned. These predictions match those of the simple model and show that, though more complex and not directly solvable, the arm reaching model exhibits the same behavior.

To extract cost ratios and proportion learned from trajectory data, we utilize a search function that minimizes the difference between the experimental trajectories and model-produced trajectories of which the cost ratios and proportion learned are known. The search function’s ability to accurately extract these values from existing trajectories is essential, thus we sought to verify its performance using an incremental approach. To verify performance, we tested the ability to fit model-generated trajectories instead of experimental data, where the true proportion learned and cost ratios are known. First, we tested a case where the search function could vary only cost weights, then we tested a case where it varied just proportion learned, finally we tested a case where it could vary both. We chose to test example trajectories near the extremes shown in Fig3A. We tested the search function’s ability to fit to combinations of two proportions learned (60% and 100%) and the high and low cost ratios (−0.0175 and −3.36, respectively).

The tests verified the search function was able to accurately recover cost ratios and proportioned learned. In the most complex case, where learning and washout trajectories were fit and both costs and learning were allowed to vary, the search function accurately captured these values across each condition, summarized in Table 1 below. The mean found proportion learned across all sets of trajectories roughly varied by only 1%, while cost ratios were slightly more variable, but within the correct order of magnitude. This also gives confidence that finding only 10 model solutions is sufficient to accurately capture the data on average. An example set of solutions for 60% and 100% proportion learned for the high cost ratio are shown in Fig 3F and 3G.

**Table 1:** Search Function Verification Results

| True Cost Ratio | True Proportion Learned | Found Cost Ratio (mean±s.d.) | Found Proportion Learned (mean±s.d.) |
| --- | --- | --- | --- |
| Low (-3.36) | 60% | -3.37±0.511 | 60.7±1.08% |
| Low (-3.36) | 100% | -3.36±0.723 | 100.7±0.860% |
| High (-0.0175) | 60% | -0.124±0.917 | 60.5±1.14% |
| High (-0.0175) | 100% | -0.0437±0.656 | 98.9±0.590% |

### Model fits to reach trajectories

With confidence that our search function can accurately find solutions, we then fit model-produced trajectories to the experimentally observed trajectories, where a single controller was used to describe all four major phases of the experiment (late baseline, early learning, late learning, and early washout), and only the model’s value for proportion learned was allowed to vary between phases. Critically, we assume that within a population, the strategy remains the same throughout the course of the experiment where each phase uses the same cost parameters.

Using this method, we found model-generated trajectories that accurately described the experimental data for both younger and older adults. The results from the best fit model for each age group are summarized in Fig 4. The model-produced trajectories are compared to the experimental data for each major phase of the experiment (late baseline, early learning, late learning, and early washout). Where the spatial trajectories are shown in Fig 4A and each of the corresponding learning metrics for each trajectory and phase are shown in Fig 4B. For both younger and older subjects, the model-generated trajectories in late learning and early washout had all learning metrics fall within the 95% confidence intervals of the experimentally observed trajectories.

**Fig 4.**
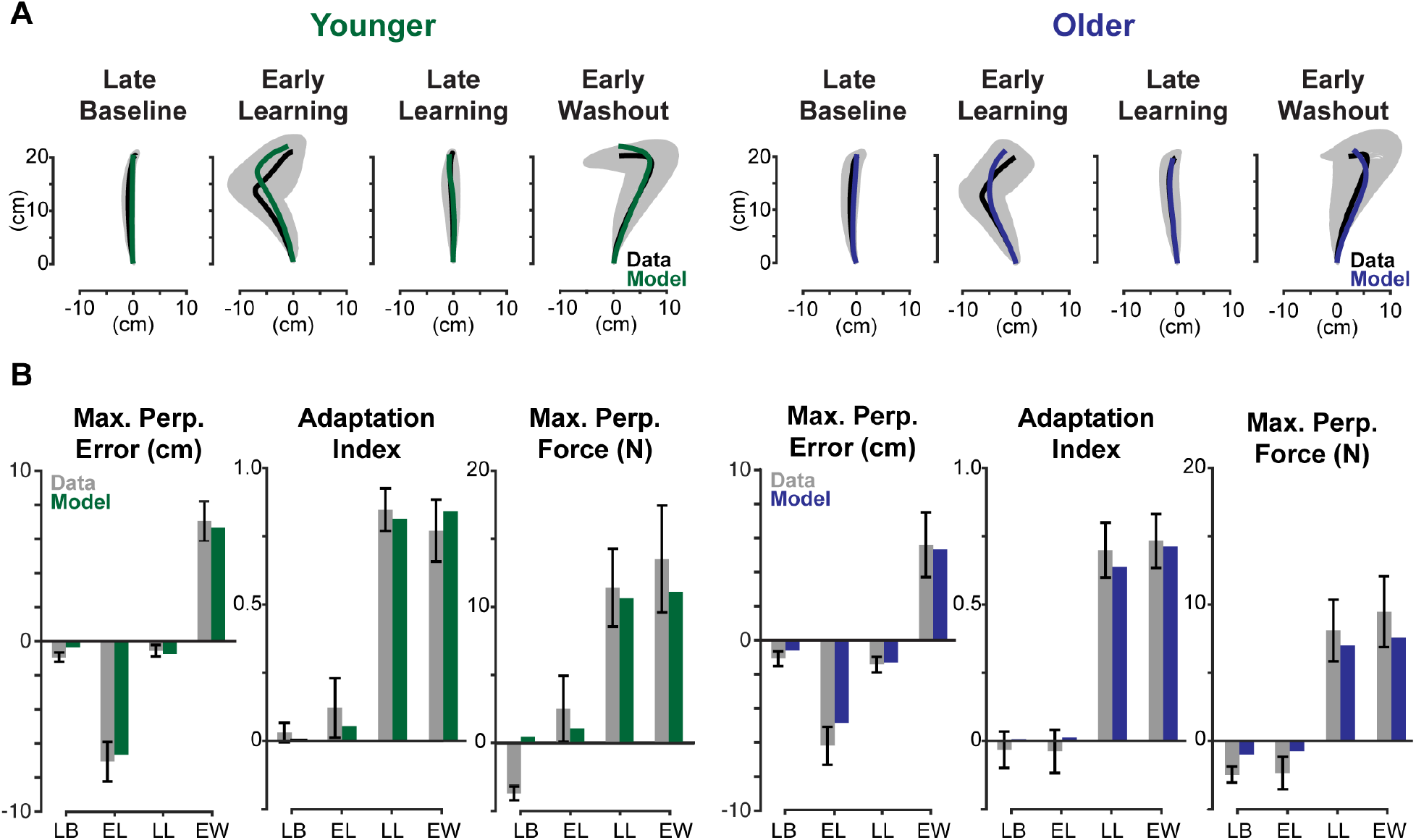
Model fits for younger and older adults. (A) Model-produced trajectories for late baseline, early learning, late learning, and early washout are compared to data from older and younger adults. Data from the experiment are shown in black with the 95% confidence interval shaded in gray. Model fits are shown in green for younger adults and blue for older adults. (B) Maximum perpendicular error, adaptation index, and maximum perpendicular force are compared for each phase. Experimental data represented in gray bars with error bars representing 95% confidence intervals, and model fits are represented in green and blue for younger and older, respectively.

Some learning metrics did not fall within the 95% confidence interval for the late baseline and early learning phases. The maximum perpendicular error and maximum perpendicular force for both older and younger adults fell outside this range. These phases, however, are exemplary in capturing the limitations of our model. Some of the natural curvature seen in the trajectories in late baseline, a symptom of biomechanical constraints and the dynamics of the robotic manipulandum, is difficult to capture with a simple point mass. These differences are small, as shown by the resulting spatial plots of the trajectories and are acceptable because our focus is on later phases of the experiment (late learning and early washout), where the model captures subject behavior more reliably. Taken together, these results demonstrate that a model that assumes subjective costs do not change over the course of the experiment can reliably capture subject behavior.

The model-based proportion learned, as previously defined, provides a latent metric for the internal model of the dynamics. When fitting all four phases, we find solutions that qualitatively match the learning process. The late baseline phase was fixed at a proportion learned of zero for both younger and older adults. We found younger adults had proportion learned values 5.3%, 79.3% and 82.3% for early learning, late learning, early washout respectively, while older adults learned 0.0565%, 76.6% and 84.7%. These findings match expectations, where, for each subject group, early learning should be close to zero and late learning and early washout should be roughly equivalent. The proportions learned found for late learning suggest that younger adults and older adults learn close to the same amount. Investigating the cost ratios which sum the kinematic costs (e.g., horizontal deviation) and divide them by the sum of effort costs (e.g., force and rate of force), we found younger adults had a log transform of the cost ratio of error to effort of −4.495, and older adults, −5.369. A more negative number, as seen in older adults, indicates a higher cost on effort relative to error. Interestingly, we see a bigger separation between cost ratios than with proportion learned in late learning. This suggests that the observed kinematic differences are due to younger adults caring more about error relative to effort than older adults, rather learning slightly more.

### Model-derived range of learning

The previous solution presents a single best fit, but we sought to determine how robust this solution was. If we assumed the proportion learned was a particular value other than those found above, could we find a set of cost weights that could adequately describe the data? If predictions from the simple model in Fig 1 are accurate, then there may be several combinations of learning and costs that could describe the experimental data. To determine the robustness of the results above, we manually derived a confidence interval for the proportion learned. We asked whether different amounts of learning could predict similar learning metrics as previously analyzed (maximum perpendicular error, maximum perpendicular force, and adaptation index). For each subject group, we re-ran the search over late learning and early washout trajectories, where we held the proportion learned constant and searched only for a set of the subjective cost weights that could describe the data. We repeated this analysis for a range of proportion learned values, from 0.4 to 1.2 in 0.05 increments.

We set a stringent set of criteria for if a model fit adequately described the data. A model fit was only deemed acceptable if the model-generated trajectories for both phases had learning metrics that fell within 95% confidence intervals (c.i.) of the real trajectory data. Statistically, those might be deemed similar, or rather, not significantly different. Using these criteria, we found several solutions that did not adequately describe the data, leaving a small window of acceptable proportion learned values for each subject group. For younger adults, adequate solutions were found with a proportion learned ranging from 65% to 90%, and for older adults, 60% to 85% of the dynamics: an overlap in proportion learned from 65% to 85%. These model fits, visualized in Fig 5, suggest that the two populations could had learned the same amount.

**Fig 5.**
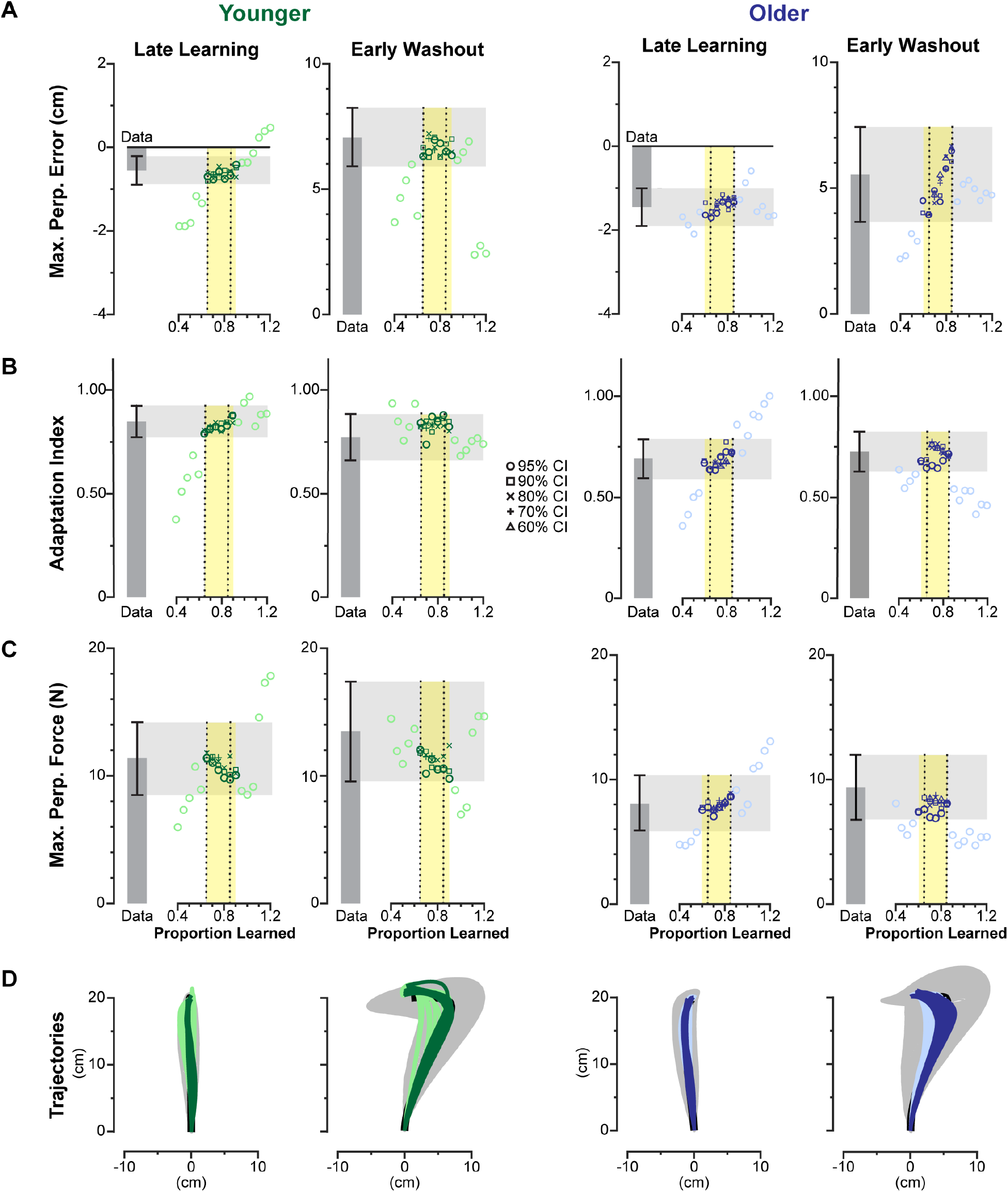
Adaptation metrics for model fits across different amounts of learning. Learning metrics from the data are shown as dark gray bars with black 95% c.i. The light gray region highlights the 95% c.i. bounds, deemed for the model to be statistically similar to the data for that phase and metric. The yellow area highlights the region that has statistically similar model fits across all three learning metrics, for both phases within a subject group (younger: green, older: blue). Each model fit is represented as a point, where acceptable model fits are highlighted in bold and unacceptable model fits are a lighter shade. Fits for each level of success criteria (95% c.i. down to 60% c.i.) are represented as different shapes per the legend. Fits for all three learning metrics (A) maximum perpendicular error, (B) adaptation index, and (C) maximum perpendicular error, and (D) each fit’s resulting trajectories are shown with the actual trajectories shown in black with 95% confidence ellipses in gray.

It is important to acknowledge that 95% c.i. gives the model a significant amount of margin to work within. We also sought to test how little margin we could give and still arrive at the same result, so we repeated the process above, but with increasingly stringent acceptability criteria. Instead of using 95% c.i., we attempted to fit trajectories across same range of proportion learned, but with learning metrics falling within the 90% c.i. (1.645 x s.e.), 80% c.i. (1.281 x s.e.), 70% c.i. (1.036 x s.e.), and 60% c.i. (0.842 x s.e.). Critically, this approach assumes that we have accurately captured the mean and are simply increasing the confidence in the data by reducing the c.i. widths. As we reduced bounds of the success criteria, the range of learning that could accurately describe the data also decreased. Shown in Fig 6, 90% c.i. had the same range of acceptable fits as 95% c.i. for both subject groups. Older adults’ most stringent successful fit was at 60% c.i., where acceptable solutions were found for 70%, 75%, and 80% proportion learned. For younger adults, the most stringent criteria where solutions were found was 70% c.i. At this level, we found solutions for 70% and 75% proportion learned. An overlap between younger and older still existed at 70% c.i., suggesting our results are robust to an increased confidence in trajectory data. At the 70% c.i. level, we see both age groups solutions converge towards 70-75% proportion learned, rather than diverge into different bins of proportion learned, further suggesting that older and younger adults may have learned the same amount. This result also corresponds well to the best-fit solution, which predicted younger adults learned 79.3% and older adults, 76.6%.

**Fig 6.**
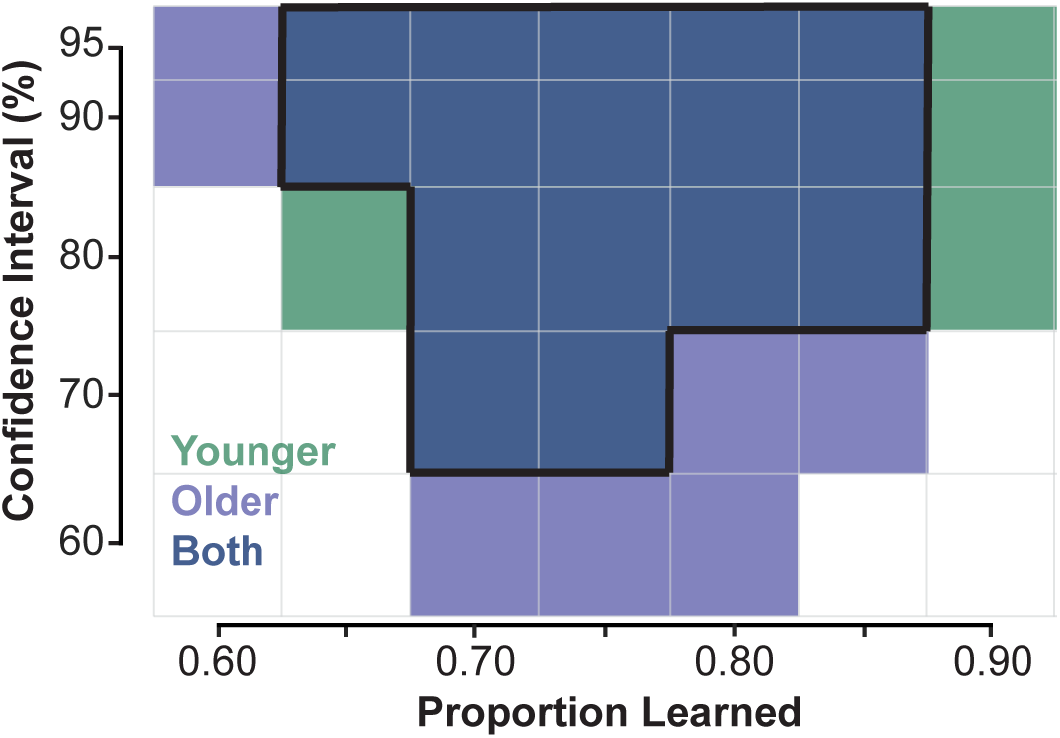
Successful model fits across different amounts of learning with increasingly stringent success criteria. Model fits with all adaptation metrics falling within the corresponding confidence interval highlighted in green (Younger), light blue (Older) and dark blue (both Younger and Older). Overlap between both subject groups is outlined in black.

This analysis has shown there are a number of solutions for both younger and older adults with equal proportions learned. If our analysis had shown there was no overlap between the proportion learned of older and younger adults, then we could posit that the two populations had learned different amounts. However, because the resulting ranges overlapped, and did so with more stringent success criteria that mimics a reduced standard error, the differences in reaching behavior between younger and older adults could be due to a difference in subjective costs, rather than a difference in ability to learn.

### The differences in subjective costs

Model fits suggest that older and younger adults may have learned the same amount, and differences in their subjective costs resulted in different reaching behaviors. We speculated whether the interaction of these subjective costs involved a consistent trade-off between effort and error, that could help define a general strategy for each population. To investigate this interaction, we used the same method to produce numerous additional model fits for both younger and older adults across their overlapping range of proportion learned (65% to 85%). We then analyzed the ratio of their kinematic costs (position and velocity terms) relative to their effort costs (force, rate of force, and control input) to investigate how these and proportion learned interact. If the predictions matched those presented in Fig 1 and Fig 3E, we would expect that greater relative costs on kinematic error could mask a deficit in learning. Similarly, we would observe that older adults, who exhibit higher error, would have consistently higher costs on effort relative to kinematic errors than younger adults.

First, we see that for equivalent trajectories within each subject group, as the proportion learned increases, the ratio of kinematic costs to effort costs decreases (Fig 7). Using a simple linear regression model to test if proportion learned significantly predicted cost ratio. The overall regression was significant for both subject groups (younger: *R^2^* = 0.0954, *F*(1,48) = 5.06, *p* = 0.0291; older: *R^2^* = 0.328, *F*(1,48) = 23.4, *p* < 0.001), where both younger and older adults have significantly negative slopes (younger: slope = −7.65, *p* = 0.0291; older: slope = −16.9, *p* < 0.001). This result matches the predictions laid out by the simple model described in Fig 1 and arm model in Fig 3E, validating that the more complex model fit to real data exhibits this same predicted behavior. Additionally, the ratio of kinematic to effort costs across this range of learning is significantly greater for younger adults than older adults, with bootstrapped 95% confidence intervals for younger: [-2.36,-1.39] and older: [-5.04,-3.87], which do not intersect. This further supports our explanation of the observed differences between younger and older adults: older adults could have learned to the same extent as younger adults; however, older adults placed a greater premium on reducing effort compared to reducing kinematic errors than younger adults.

**Fig 7.**
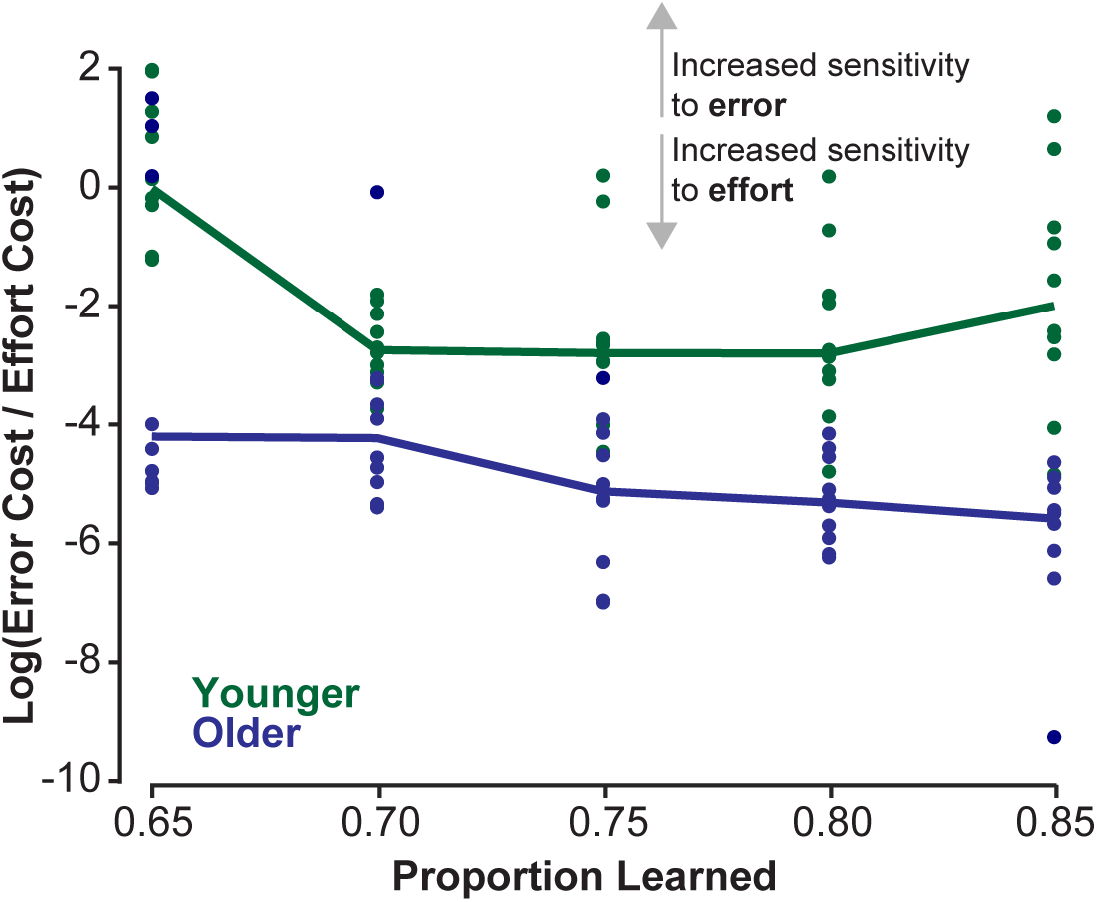
Older adults have a lower cost on kinematics relative to effort than younger adults. The ratio of kinematic costs to effort costs are shown for younger (green) and older (blue) adults. Each dot represents a model fit that has learning metrics which fall within 95% c.i. of the experimental data for that subject group for the prescribed proportion learned. Solid lines connect the mean cost ratio for each proportion learned per subject group. A higher ratio of kinematic costs to effort costs (larger value in the y-axis) can be interpreted as having an increased sensitivity to error, where a lower cost ratio could be interpreted as an increased sensitivity to effort.

### Analyzing individual performance

Another approach to investigating strategic differences is to extract subjective costs and proportions learned for each individual, then collect and compare across the subject groups. For each subject, the search function varied cost weights and a single proportion learned value that was the same for each phase. Two example subjects, one younger and one older, are shown in Fig 8A, where we see the model is able to accurately describe these subjects’ trajectories. One younger adult did not have an acceptable trajectory for late learning, thus was left out of the analysis. One younger adult and one older adult did not have acceptable model fits and were left out of the statistical comparisons between groups. Compiling the findings from each individual fit, we find younger adults learned slightly more on average than older adults (mean±s.e., younger: 81.9±4.87% older: 74.7±5.40% *t*(23) = 0.975, *p* = 0.340, Fig 8B), though the groups are not significantly different. Additionally, we find that younger adults cost ratio is slightly higher (−4.51±0.281 versus −4.99±0.225, *t*(23) = 1.34 *p* = 0.193, Fig 8C) though it is also not significantly different between younger and older adults.

**Fig 8.**
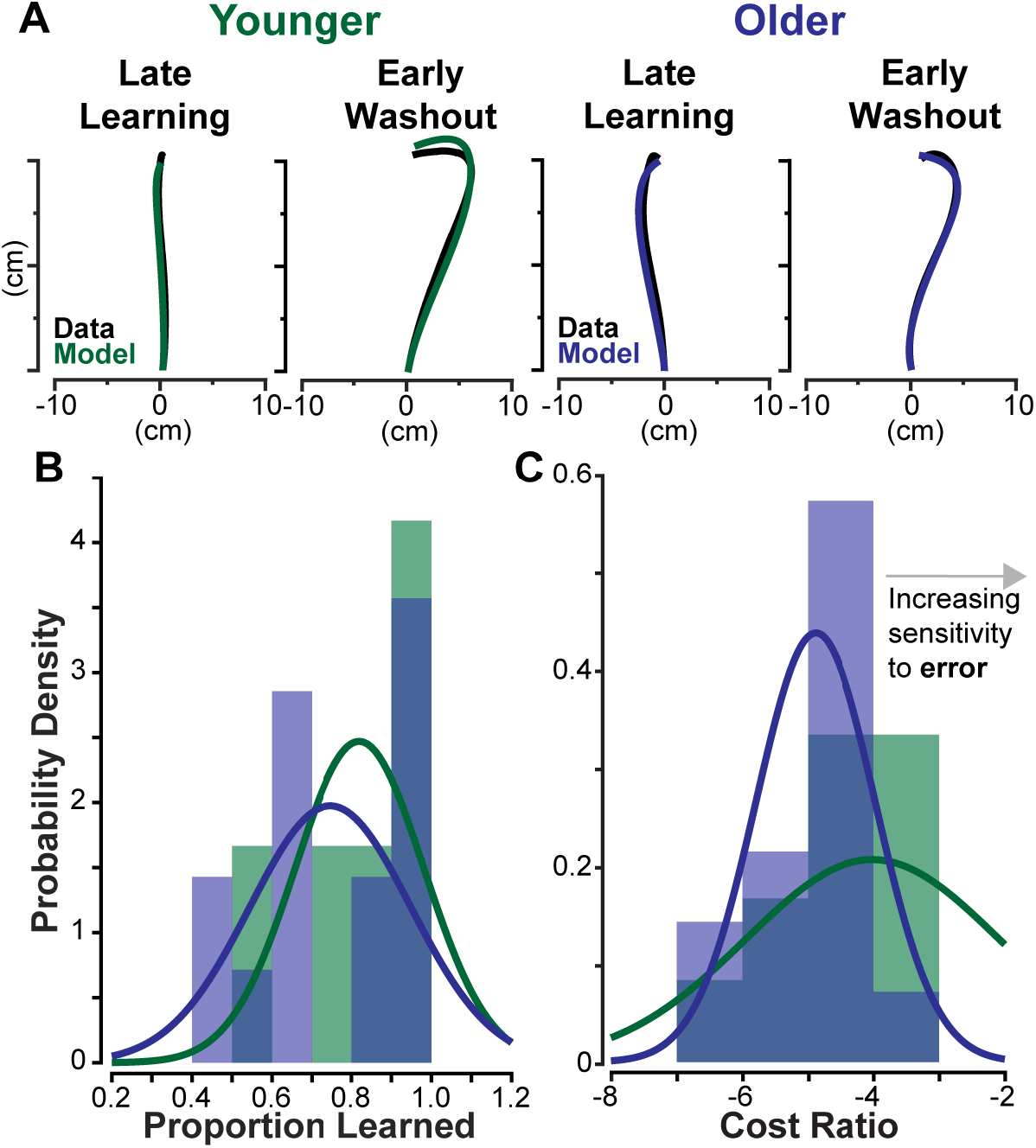
Individual Subject Trajectory Fits. For each subject, trajectories were fit to late learning and early washout phases. (A) An example fit for a younger subject is shown on the left and for an older subject on the right. The original data is shown in black, while the model produced trajectories are shown in green and blue. (B) The distribution of proportion learned for each subject is plotted, where data for younger adults are shown in green, and older adults in blue. A normal distribution was fit to each dataset and is overlayed in bold. (C) Follows the same labeling and color scheme as (B) and shows the distribution of cost ratios for each subject group, where less negative values demonstrate an increased weight on error costs relative to effort.

Due to the small sample size and limited number of trials, the individual analysis is limited in its utility for this particular dataset. While it does not find any statistical differences between groups, it weakly suggests that both learning and cost ratios may differ, with younger adults learning more and having a higher cost on kinematic error than their older counterparts. Moreover, the average proportion learned and average cost ratio for each group roughly matches the values we obtained through the group analysis. At a minimum, analyzing individuals offers an alternative framework to determine subjective costs and learning and enable within subject comparisons. However, for this particular dataset, the group-averaged approach seems more appropriate in order to avoid issues with losing data.

Collectively, our analyses suggest that younger adults could have different subjective costs than older adults, and this may be the cause for observed kinematic differences. When analyzing averaged trajectories for each group, we found similar proportioned learned between younger and older adults (79.3% and 76.6%), but different cost ratios (−4.495 and −5.369). This was further supported by investigating the range of learning that could accurately describe the learning and washout phases for each group, resulting in significant overlap between subject groups (65% to 85%). Further, when the bounds for an acceptable fit became more stringent, both groups converged to the same amount of learning (70-75%). Investigating the set solutions for each subject group, across all values of proportion learned, younger adults had a significantly higher cost ratio of error relative to effort. Finally, when analyzing individual subjects, we see again that proportion learned did not significantly differ between younger and older adults, though this may be due to the small sample size and lack of power. However, the values found for both proportioned learned and cost ratios mirror analyses performed on the group average data quite well. This suggests that the kinematic differences observed between younger and older adults may be a result of different sensitivities to effort expenditure and error, rather than a difference in learning.

## Discussion

Our analysis suggests that larger kinematic errors do not necessarily imply less learning, and that we must consider subjective strategies when assessing learning. Using our model, we find that older and younger adults reaching trajectories can be explained with similar amounts of learning, despite their large kinematic differences. These differences in motor behavior may be attributed to older adults caring more about effort relative to kinematic error than younger adults. Our model assumes that objective effort is similar in younger and older adults, which is consistent with measured metabolic power in Huang and Ahmed 2014 [29]; thus, our results suggest that it is the subjective weighting of effort that differs. [48]. An alternative explanation is that the objective effort costs are higher in older adults compared to younger adults. Locomotion studies have shown older adults incur greater metabolic cost, an objective measure of effort, compared to younger adults [49,50]. While metabolic cost has been measured in both younger and older adults performing motor learning tasks, we await a direct comparison of the metabolic cost between groups. Our findings, in their current form, cannot distinguish between a higher subjective effort cost versus a higher objective effort cost in older adults.

Additionally, our results question the validity of adaptation index used in Huang and Ahmed 2014 [29] as a measure of learning in curl field experiments. We show that the same trajectory can be produced across different proportions learned, and because the adaptation index is calculated using trajectory data, the adaptation index will not correlate to the model-based, latent state of proportion learned. This suggests that adaptation index may be a poor indicator of learning between groups, as it inherently assumes a single strategy where kinematic error-canceling is more highly weighted than effort.

Of note, our finding may be unique to the force-field adaptation paradigm. Using visuomotor rotations may eliminate any effort-centric strategic differences. However, visuomotor rotation experiments probe different mechanisms in the motor learning domain. Force-field adaptation tasks use proprioceptive feedback to make online and trial-to-trial corrections, while visuomotor rotations use visual information as a feedback signal. Because force-field adaptation tasks probe both effort and error simultaneously, the paradigm may better emulate motor tasks encountered in the real world.

Additionally, the significant difference in learning metrics between older and younger adults may be unique to the protocol used for this force-field adaptation task. The Huang and Ahmed 2014 study employed a protocol that may have put an added premium on the effort of the task [29]. Due to the desire to collect metabolic data, subjects reached nearly continuously in both the anterior and posterior directions during each trial block. Other studies that did not find a difference between age groups used different reach distances, reach time constraints, curl field strengths, reach directions, and/or inter-trial intervals [41,43,44]. All these nuanced differences in protocols could have increased the effect observed by Huang and Ahmed [29].

The data from Huang and Ahmed [29], while a unique finding in geriatric research, does have some drawbacks due to its limited sample size. Though the sample size was sufficient to find statistically significant difference in learning metrics, the confidence bounds combined with the degrees of freedom of the model may provide enough flexibility so that an overlap in learning would necessarily exist. An increased sample size would be helpful in drawing stronger conclusions. This shortcoming is especially present in the individual analysis, where our objective function was necessarily different due to only some of the subjects experiencing a channel trial in the early washout phase. With a different protocol or an increased sample size, results in comparing between older and younger adults may be more conclusive than what is offered in this analysis. Nevertheless, this example dataset shows how these methods can be applied to extract latent variables and offer an alternative explanation to assumed differences between age groups.

It is worth acknowledging that our model was purposefully simple, so differences between subject groups are more easily interpretable. However, as with all modeling studies, there is a trade-off between biological realism and model complexity. It could be the case that a different, or more realistic model would result in different findings. A higher fidelity model, reducing the influence of underlying assumptions, could be accomplished through a few different means.

The first improvement lies in the dynamical model. For instance, the use of a non-linear model of the arm could offer improvements over a point mass. As already discussed, this could improve the fits in the late baseline phase. This type of model could exhibit some natural deviation from the centerline and reduce costs penalizing non-straight reaches. Additionally, a more complex model that allows for co-contraction could also offer further insight into effort costs and error-canceling strategies that are independent of the perturbation. Our model, only considers motor commands as a basis of effort, but subjects could be exerting more effort through co-contraction that is unable to be captured by our model. Another improvement could be to include off-diagonal terms in our cost functions to capture interaction terms between state variables. This would improve the quality of our fits, but in turn, make the model prone to over-fitting and make the subjective costs less interpretable. Finally, developing a model that incorporates motor noise, sensory uncertainty or delay may offer an alternative explanation to the differences in behavior [51]. This could be a driving force in their movement strategy that could be reflected in learning rate or trajectory differences, and potentially mask differences in learning or subjective costs. Overall, these improvements to the model could help tease out the specific differences between subject groups, and ultimately determine whether populations are learning less or compensating in a different manner.

While it is important to extract certain behaviors from observed trajectories such as arm reaches, using metrics such as maximal values cannot account for the temporal, stochastic, and highly dynamic factors surrounding human motor control. Strong conclusions about how two populations learn differently should be extremely thorough and consider multiple metrics. The powerful framework of optimal control enables us to compare temporal data to temporal data, extract valuable information from these models, and estimate the hidden value of how much a person has learned. Deducing whether a difference in behavior is due to a difference in learning, a difference in subjective costs, or a combination of the two is still unanswered; however, we offer a framework which can probe these differences.

Our results show that subjective movement strategies can mask the latent variable of how much a person has learned. Additionally, we have shown that both older and younger adults adapt their reaches to a curl field, but whether they definitively learn to different extents remains unclear. If younger and older adults learn to the same extent, our model offers a plausible explanation that behavior differences between older and younger adults are caused by older adults caring more about effort relative to kinematic errors. We show that using learning metrics alone gives insufficient insight into the adaptation process. In future studies investigating how much a person or population has learned, it is imperative to consider their implicit strategies.

## Materials and Methods

### Experimental Setup

This experiment used data from Huang and Ahmed 2014, which investigated differences in learning between older and younger adults [29]. We will briefly review the experiment here and refer the reader to the original publication for greater detail. Huang and Ahmed’s study [29] utilized data from eleven older adults (mean±s.d., age 73.8±5.6 years) and 15 younger adults (23.8±4.7 years). The analyses in this study uses data from 15 older adults (age 72.9±5.44, 7 female, 8 male) and 13 younger adults (age 24.7±4.6, 13 female, 2 male). In the original submission, four older adults were not included due to poor quality of metabolic data. However, because this study analyzed reaching trajectories only, these subjects’ data could be included. Additionally, two younger adult subjects’ data were not used due to corrupted or inaccessible reach trajectory data. Despite these different subject numbers, conclusions from the original manuscript are the same.

Each subject made targeted reaching movements while grasping the handle of a robotic arm. The experimental setup is visualized in Fig 2A. Subjects were seated with their right forearm cradled and the computer screen set at eye level. Reaches were made in the anterior and posterior directions. Successful reaches needed to reach the within a movement time between 300 and 600 ms. Fig 2B outlines the various dynamics subjects experienced throughout the course of the experiment. Each subject made 900 reaches with the robotic arm, 450 anteriorly, 450 posteriorly. For the middle 500 reaches, subjects were exposed to a velocity-dependent force (curl) field. The forces imparted by the robotic arm per Equation 5, where the forces, *f_x_* and *f_y_,* imparted on the hand were proportional and perpendicular to the velocity of the hand, *v_x_* and *v_y_,* scaled by the curl field gain, *b*. In this experiment, *b* is −20 N-s/m. The progression of trials was broken into three blocks: 200 trials with no forces (baseline), 500 trials with the curl field on (learning), and a final 200 trials with no forces (washout).

Throughout the 900 trials, subjects were exposed to one force channel trial every five trials, pseudo-randomly dispersed. The channel trial forced subjects to reach in a straight line, while simultaneously measuring the forces exerted on the robotic arm. The robot arm enforced the channel trial using a horizontal force relative to the horizontal position and velocity, summarized in the equation below:

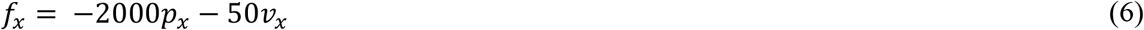

As subjects adapted their reaches to the curl field, they began to anticipate the perturbing forces. On channel trials, subjects’ anticipatory compensation resulted in exerting force against the channel, opposite the direction of the curl field perturbation. The measured force trace, often analyzed in conjunction with the velocity trace, provides insight into how well subjects have learned the novel dynamics. The exact process of estimating this value is detailed in the *Learning Metrics* section below.

### Data Preparation

#### Trajectory Data

To analyze the performance of our model, we considered only the outward reaches from the final ten trials in the late baseline phase and late learning phase, and the outward reaches from the first five trials of the early learning and early washout phase. All position, velocity and force data were collected at 200 Hz. To be included in the analysis, the subject must have reached the target between 250 ms (50 samples) and 1500 ms (300 samples) after the target appeared. Movement onset was defined as when the anterior velocity was ≥ 0.03 m/s towards the target, and movement termination was defined as when the cursor was within the target area and anterior velocity was ≤ 0.03 m/s. If the trial never reached the target area, and/or did not sufficiently slow down, it was still included, but subjected to the reach time criteria mentioned above. Collectively across experimental phases and subject groups, 96% of trials met these criteria. Younger adults had 5.8% of trials that did not meet the time criteria and older adults, 2.9%. Of the failed trials, 74% were in the Early Learning phase, and 21% in the Early Washout phase. Finally, if data were corrupted or the robotic arm setup experienced any issues, these trials were thrown out (<0.1% of trials). The trials that met these criteria were averaged across each subject with each resampled to the mean trial length of that subject’s trajectories. This set of trajectories was used for fitting individual subject data. To obtain group averages for the younger and older subject groups, the mean trial length across subjects was calculated, each subject’s mean trajectory was normalized to that trial length, then averaged across subjects. Channel trials were considered separately from the non-channel trials but used the same criteria and processing method. Due to the pseudo-randomly dispersed trials in the original experiment and the criteria set for an acceptable reach, there was typically one or zero channel trials for a subject in the set of trials considered for each phase. However, it was essential to only consider the first five reaches in early adaptation and early washout, because adaptation and de-adaptation occur very quickly. We chose these stricter inclusion criteria to represent model assumptions more accurately, thus performance metric values presented here are slightly different than the numbers in the original manuscript; however, they do not alter the conclusions made in the previous manuscript.

#### Learning Metrics

From the time-normalized average trajectories, we calculated commonly used learning metrics: maximum perpendicular error, maximum perpendicular force, and adaptation index. Maximum perpendicular error is calculated as the largest absolute perpendicular deviation from a theoretical straight line that connects the start position to the target position. Often, the net perpendicular deviation is considered, where the average perpendicular deviation in the training phase, or baseline phase, is subtracted from the perpendicular error in the novel environment. Using the net deviation accounts for any natural curvature in reaches due to the biomechanical constraints of the arm and allows for a better within-subject analyses. In this experiment, however, we were more concerned about the comparison between subject groups, so total perpendicular error was a more appropriate metric.

Because this experiment was a force-adaptation task, it was also useful to consider how the anticipatory force changed over the course of the experiment. Using data collected from channel trials, we calculated the maximum force that each subject pushed against the channel, perpendicular to the direction of movement. A positive value indicates a force in the right-hand direction against the perturbing force, while a negative value indicates a force in the left-hand direction with the perturbing force. Accordingly, this value can be compared to an expected maximum perpendicular force, calculated from the maximum velocity and curl field gain.

A more comprehensive metric for adaptation, assuming the strategy is to reach as straight as possible, is to compare the entire force trace to the entire velocity trace in a channel trial. In a curl field trial, the horizontal force is applied to the hand proportional to the vertical velocity through the scalar value, *b,* as per Equation 4. If a trajectory accurately anticipates and compensates for these forces, the force trace measured in a channel will be approximately equal to *b* times the velocity profile. If the dynamics are underestimated, the force profile will be equal to the velocity profile scaled by some value less than *b*. Thus, using the measured horizontal force trace and dividing it by the vertical velocity trace, the scalar value that best approximates this linear relationship is an estimate for the amount of adaptation that has occurred, which we call the adaptation index.

#### Arm Reach Model

We employed a discrete-time, two-dimensional, finite-horizon, linear quadratic optimal control model using a symmetrical point mass to describe the arm reaching trajectories. The model included an internal estimate of the state dynamics to calculate the control law (referred to as the proportion learned, above), where all states were deterministically observable and there was no system variability or uncertainty. The system dynamics are governed by the following equation

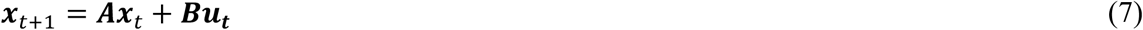

where ***A*** and ***B*** are the dynamics of the system, and ***x*** is the state vector defined as follows:

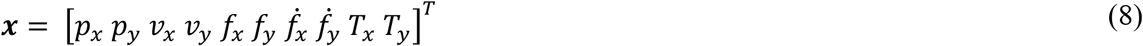

The variables *p_x_* and *p_y_* represent hand position, *v_x_* and *v_y_,* hand velocities, *f_x_* and *f_y_* are hand forces, *ḟ_x_* and *ḟ_y_* are the rate of change of hand forces, and *T_x_* and *T_y_* are target position. In this paradigm, target position is constant. The motor commands, *u_x_* and *u_y_* are the second derivative of force. The matrices ***A*** and ***B*** encapsulate the dynamics of the curl field and channel trials, as outlined in Equations 5 and 6. Additionally, a separate, but similar matrix, ***Â***, represents the internal model of the dynamics, which includes a subject’s estimate of the curl field gain instead of the true value.

The cost function used to calculate the control law is defined as:

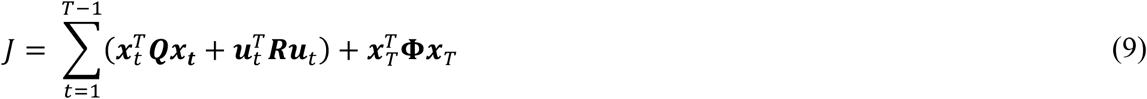

where the matrices ***Q**, **R***, and ***Φ*** are symmetric matrices, which penalize state tracking, control input and terminal state, respectively. The matrices represent these penalties as cost weights along the diagonal of each matrix. Within ***Q***, there are separate costs for horizontal deviation from target/centerline ((***Q***(1,1) = ***Q***(9,9) = −***Q***(1,9) = −***Q***(9,1)), vertical distance from target (***Q***(2,2) = ***Q***(10,10) = −***Q***(2,10) = −***Q***(10,2)), horizontal velocity (***Q***(4,4)), force (***Q***(5,5) = ***Q***(6,6)), and rate of force (***Q***(7,7) = ***Q***(8,8)). Within ***R***, there is a cost on the second derivative of force (***R***(1,1) = ***R***(2,2)). And within ***Φ,*** there are costs on the end position relative to target position (***Φ***(1,1) = ***Φ***(2,2) = ***Φ***(9,9) = ***Φ***(10,10) = −***Φ***(1,9) = −***Φ***(9,1) = −***Φ***(2,10) = −***Φ***(10,2)), the end velocity relative to zero (***Φ***(3,3) = ***Φ***(4,4)), end force relative to zero (***Φ***(5,5) = ***Φ***(6,6)), and end rate of force relative to zero (***Φ***(7,7) = ***Φ***(8,8)). All other elements in these matrices were zero. Collectively, these matrices represent the subjective costs for a movement and are and are used to formulate a movement plan where the control sequence, determined by the control law, is calculated with these cost matrices and the estimated dynamics, ***Â*** and ***B***. Trajectories were simulated by running the control sequence forward in time through the true dynamics (either the with or without a curl field or within a force channel).

### Trajectory Matching

When fitting the model to the experimental results, we approached the trajectory matching problem using optimization techniques. We sought to minimize an objective function that described the data through varying the cost weights in ***Q, R**,* and ***Φ*** used to calculate the optimal control model and trajectory. In some cases, the internal model of the dynamics (i.e., proportion learned), was allowed to vary as well. We used MATLAB’s constrained minimization function, *fmincon,* designed for nonlinear optimization problems. The minimizing solution was obtained by comparing results from multiple restarts with randomized initial parameter values. Of the multiple restarts, the solution that resulted in the smallest value of the objective function was chosen.

### Objective Function

Our initial analysis of group data considered four phases of the experiment: late baseline, early learning, late learning and early washout, and the fit sensitivity and cost ratio analyses considered only late learning and early washout. Within each phase, both channel trials and non-channel trials were considered in the objective function. Within the objective function, the weighted sum of the z-scores of the data’s end point, maximum perpendicular position (error), adaptation index, and maximum perpendicular force are penalized. When analyzing trajectories without confidence intervals, as is the case with the model validation and individual data, the objective function penalized these same metrics, but did not z-score them.

To find the trajectory that minimizes the objective function, the proportion learned varied in some cases, and the weights on each cost parameter were varied. Cost parameters that penalized the horizontal hand position, vertical hand position, hand force, rate of change of hand force, second derivative of hand force (control input), terminal position, terminal velocity, terminal force, and terminal rate of change of force were allowed to vary. Each of these parameters were along the diagonal of cost matrices ***Q***, ***R***, and ***Φ***. Constraining these matrices to be diagonal limited the quality of fits thus our solutions represent a lower-bound on the quality of fits achievable. In the initial fit across all four phases, the value of the internal model’s proportion learned for early learning, late learning and early washout were allowed to vary in finding the best fit, while late baseline was fixed at a value of zero (i.e., no learning). In the subsequent analyses of group data, we find how sensitive the best fit trajectory was, the internal model of the dynamics was fixed at values from 40 to 120% of the curl field gain in the late learning phase, while the other parameters were still allowed to vary. Ultimately, this produced a range of learning values and subjective costs that accurately describe the data, where the learning metrics must fall within the set maximum confidence interval (initially 95%). The intersection of proportion learned between subject groups was investigated further using the same method described above, but with more stringent success criteria. When analyzing individual subject data, proportion learned and cost weights could vary, and the objective function did not z-score the learning metrics.

### Search Function Performance

We took an incremental approach to verifying search function’s performance, where the search function could vary cost weights, proportion learned, or both, to fit a set model-generated trajectories. We tested the search function’s ability to fit to combinations of two proportions learned (60% and 100%) and the high and low cost ratios (−0.0175 and −3.36, respectively). Initially, we fit only a single trajectory: the “learning” phase reach in the curl field, then we tested the search function with two trajectories: the “learning” phase in the curl field and “washout” phase in the null field. The latter, with two trajectories, is what we use throughout the paper except for the initial fit which considered all four major phases. Within each category we performed the search in three ways: holding costs constant and searching for the proportion learned, holding proportion learned constant and searching for costs, and finally letting both costs and proportion learned vary. For each search, we found ten best fit solutions, then calculated the mean cost ratio and proportion learned across these solutions.

### Model Sensitivity Analysis

Simulated reaches whose learning metrics fell within the specified confidence intervals (initially 95%) of the experimentally obtained metrics were considered statistically indistinguishable from the data. Only solutions that had all learning metrics that fell within this range for both late learning and early washout were considered acceptable solutions. After these solutions were found, they were checked by comparing their spatial plots to that of the data.

First, we compared the resulting ranges of acceptable values of proportion learned with 95% c.i. If there was no overlap, then we could conclude the two populations had learned a different amount. If there was overlap, then investigating how the specific cost parameters differ between subject groups could offer an explanation for the differences in reaching behavior. Next, we tested the robustness of this result by enforcing more stringent success criteria. Following the same procedure, a solution was only deemed acceptable if the learning metrics fell within an incrementally decreasing confidence bound. We chose to investigate 90% c.i., 80% c.i., 70% c.i, and 60% c.i. These bounds corresponded to incrementally decreasing the acceptable range of learning metric values. As with the previous, only a solution that had learning metrics fall within these bounds were deemed acceptable.

To more easily interpret the motor strategy differences, individual costs were categorized and combined into two types: kinematic or effort costs. Kinematic costs included costs on position (perpendicular error, distance to target, end position). Effort costs included force and the derivatives of force states (*f_x_, f_y_, ḟ_x_, ḟ_y_, **u**).* Costs for each state were normalized by the sum of the squared states for to account for the difference in number of samples between subject groups. Each category of costs was summed together, then to account for a potential uniform scaling of cost weights, normalized kinematic cost is divided by the normalized effort cost to create a metric that encapsulated each group’s subjective value of kinematic versus effort costs, referred to as the cost ratio. For each subject group, the log transform of the set of all cost weight ratios for all proportions learned were sampled with replacement 10,000 times to provide the estimated 95% confidence intervals.

### Fitting to Individual Subjects

To analyze trajectories for individual subjects, we used the same inclusion criteria as the averaged group data. The same windows of trials were used: the last ten reaches of the late learning phase and first five reaches of early washout. Anterior reach trajectories were averaged across these windows to provide a single trajectory for late learning and a single trajectory for early washout per subject. In taking this approach, data is more sparse, where some individuals have only one or no successful catch trials for a given phase, which are essential for calculating maximum perpendicular force and adaptation index. Rather than not consider a number of subjects due to lack of data, and to be consistent across how we fit all subjects’ trajectories, we use a more constrained objective function that does not z-score the learning metrics and only penalized deviation from the mean trajectories’ maximum perpendicular error and end point accuracy. The search function was allowed to vary cost weights and a single value proportion learned that was used for both late learning and early washout to find the best fit solution. Once a solution was found that met the maximum allowable objective function, we combined the cost ratios and proportions learned across subjects, then compared distributions between younger and older adults.

## Acknowledgements

A special thanks to Helen Huang for sharing her dataset.

## Notes

https://github.com/neuromechanics-at-cu/learning_vs_minding

### Competing Interest Statement

The authors have declared no competing interest.

### Summary of Updates

Added discussion on limitations due to sample size. Revised individual subject analysis.

